# Scalable tagging of endogenous genes by homology-independent intron targeting

**DOI:** 10.1101/412445

**Authors:** Yevgeniy V Serebrenik, Stephanie E Sansbury, Saranya Santhosh Kumar, Jorge Henao-Mejia, Ophir Shalem

## Abstract

Genome editing tools have simplified the generation of knock-in gene fusions, yet the requirement for gene-specific homology directed repair (HDR) templates still hinders the scalability of most approaches. Here, we combine intron-based protein trapping with homology independent repair-based editing and demonstrate precise and efficient gene tagging that can be easily scaled due to use of a generic donor. As editing is done in introns, this approach tolerates mutations in the unedited allele, disruptive indels, and allows for flexible donor and sgRNA design.

## Main

Fusing endogenous proteins with fluorescence or epitope tags is a widely used and essential approach for studying proteins within their natural regulatory context. The advent of CRISPR tools for modifying the genome^1-3^ has made this easier and even more accessible, yet scalability is still very limited. The need for a gene-specific homology directed repair (HDR) template requires costly synthesis or labor-intensive molecular cloning, and since precise targeting must be achieved in frame with the coding sequence, it necessitates careful design of reagents and screening of clonal cell lines to avoid disruptive editing at the non-tagged allele. A recent development of split fluorescent proteins has simplified the generation of fluorescent fusions, since only a minimal tag is required for genomic knock-in^4-6^. Nevertheless, these endogenous tagging methods still require individual HDR donors. Several approaches to develop generic exon-tagging methods have been demonstrated^7,8^, but because these require precise tagging at the coding sequence, they are limited in design flexibility and are prone to disruptive mutations at the non-tagged allele, as well as to indels within the tagged allele that can lead to frameshifts. Derivative strategies have been developed to increase the efficiency of homology independent repair-dependent tagging methods but at the cost of no longer utilizing a generic donor^9^.

An alternative approach for generating endogenous fusions is by random integration of synthetic exons delivered by transposons or retroviral particles^10^. This approach, known as “protein trapping” or “CD-tagging”^11^, is not restricted to small donors and has been used in both model organisms^12-14^ and mammalian cells^15,16^. While protein trapping is inexpensive and scalable, the random nature of tag integration precludes its use for the generation of curated libraries of fusion cell lines.

Here, we combine homology independent repair-based editing^9^ with the use of a generic synthetic exon donor containing a fluorescence tag to perform targeted protein trapping at intronic locations (Figure 1A). This approach is efficient, easy to implement, and does not limit the size of the donor. Furthermore, in contrast to generic exon tagging, generic intron tagging benefits from increased flexibility of the donor design enabled by the splice acceptor and donor sites: any incorporated vector sequence external to those sites has no effect on the coding sequence of the tagged protein. Generic intron tagging also uniquely tolerates mutations in the non-tagged allele, as those are intronic and typically non-disruptive, as well as indels that flank the inserted donor as a result of editing that could lead to frameshifts in an exonic setting. Because the donor is generic, the generation of additional fusion cell lines only requires the cloning of additional intron-targeting sgRNAs.

**Figure 1:**
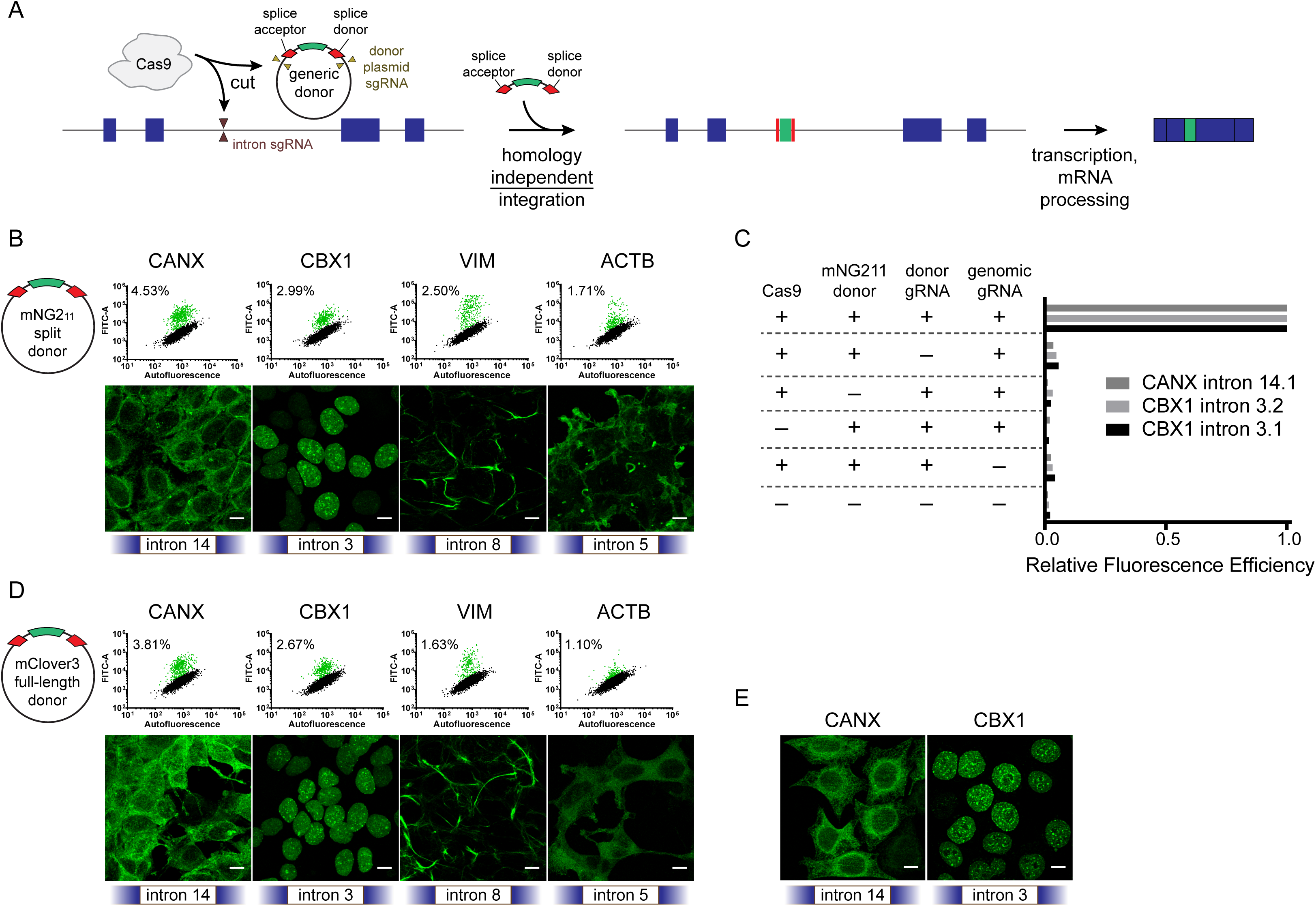
Homology-independent generic intron tagging enables efficient and easy generation of endogenous fusions. (**A**) Illustration of the tagging approach: Double-strand breaks are generated in the intron and donor resulting in the addition of a synthetic intron and fusion of the tag to the coding sequence. (**B**) Using a small donor composed of the mNG2_11_ epitope flanked by splice acceptor and donor sites results in efficient tagging of *CANX, CBX1, VIM*, and *ACTB* at the indicated introns (all sgRNA target 1), as observed by flow cytometry (upper panels) and by confocal microscopy (lower panels). Percentages in the dot plots represent the green population as a subset of the total. (**C**) All transfection mix components are required for tagging of *CANX* intron 14, sgRNA target 1 (14.1), and of *CBX1* intron 3, targets 1 and 2 (3.1 and 3.2). The table indicates which component was removed and bar plots represents the relative percentage of fluorescence-positive cells compared to the full mix. (**D**) Tagging using a full-length mClover3 fluorophore as a donor. Note that a difference in localization is observed for *ACTB* between the small and large tag. (**E**) Tagging of *CANX* and *CBX1* in HeLa cells. All images are maximum projections of z-stacks and scale bars correspond to 10 µm.

Initially, we designed a plasmid donor that contained the mNG2_11_ tag, part of a recently-published split fluorophore system^4^, flanked by linker sequences and splice acceptor (SA) and donor (SD) sites (Supplementary Data 1). We embedded this sequence between two identical sgRNA target sites chosen to have minimal off-target activity in the human genome, such that cutting of the plasmid in cells generates a linear DNA donor molecule. Plasmids containing SpCas9, sgRNAs against the donor plasmid, the donor plasmid itself, and intron-targeting sgRNAs were transfected into 293 cells already stably expressing mNG2_1-10_. Proteins successfully tagged with mNG2_11_ emit a fluorescence signal upon binding of mNG2_1-10_. Using this approach, we were able to tag four tested genes with well-established localization patterns, *CANX, CBX1, VIM* and *ACTB*, at a frequency that enabled easy isolation of both clonal and polyclonal tagged populations of genes (Figure 1B). To test that our tagging approach was indeed mediated by double-strand breaks in both the genomic sequence and the donor plasmid, we removed each individual component of the transfection mix and found that efficient tagging indeed required all components (Figure 1C). We then tested the feasibility of integrating larger donors by replacing the mNG2_11_ epitope with a full mClover3 fluorescence protein (FP) and found comparable integration efficiencies (Figure 1D). Interestingly, integrating a full-length FP in *ACTB* resulted in a lower expression level and a diffuse localization pattern, consistent with the production of non-functional protein (right most panels in Figure 1B and 1D). Lastly, to verify that the activity we observed was not specific to 293 cells, we also successfully tagged HeLa cells (Figure 1E).

Unsuccessful tagging can be a result of inefficient sgRNA cutting, low donor integration, inefficient splicing, or a fusion location that disrupts protein expression. To start investigating these alternatives, we chose two genes, *ACTB* and *CANX*, and designed nine sgRNAs for each that spanned three introns. We then measured tagging efficiency and the expression levels of tagged cells for each of these locations (Figure 2A,B). We found that both tagging efficiency and expression levels roughly correlated within the same introns, indicating that the location of the fusion is a more critical parameter than the choice of the sgRNA within an intron. We further validated this by testing for donor integration in the total transfected cell populations using pairs of primers that were designed to amplify the genomic DNA to donor junction on both sides of the donor (Figure 2C, diagram). We were able to identify integration for all sites (Figure 2C), with no discernable directional preference (Figure 2C, lower panel). Tandem insertions were also prevalent as evidenced by the upper bands that correspond to twice and sometimes three times the expected molecular weight. We also used Sanger sequencing on some of the amplified junctions and found accurate integration sometimes flanked by junction indels (Figure 2D), further emphasizing the advantages of targeting introns using such an approach.

**Figure 2:**
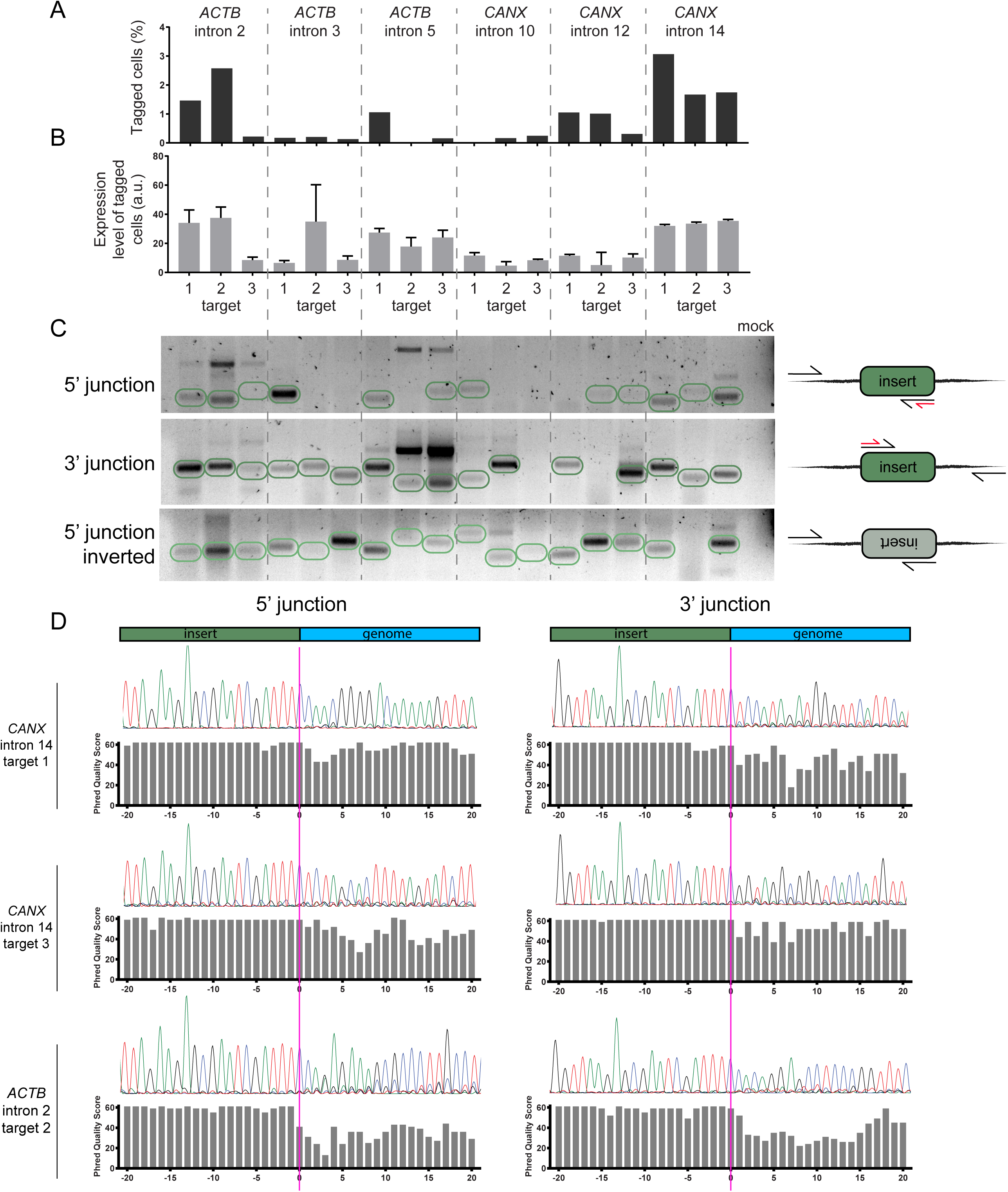
Successful tagging is mostly determined by the choice of intron. **A**) Tagging with mNG2_11_ across introns in *ACTB* and *CANX*. Bar plots represent the percent of fluorescence-positive cells for each sgRNA position. (**B**) Expression mean and standard error for positive cells in each location. Sample sizes are proportional to the bar plots in part A. (**C**) Gel image showing the amplification of donor to genomic DNA junctions, as illustrated in the right-hand diagrams. Expected band sizes for insertion of a single copy of donor are circled in green. In the diagrams, black arrows represent primer sites for amplification and red arrows represent primer sites for sequencing in part D. The last lane corresponds to a PCR reaction with primers for *CANX* intron 14, target 1, but without a template. (**D**) Sanger sequencing of donor to genomic DNA junctions shows de-dephasing at the donor and genomic DNA junction which indicates indels at the integration site.

To summarize, we show that generating endogenous fusions by generic intron tagging is efficient and easy to implement. As editing is done in introns, this approach tolerates indels both in the untagged allele and in the sequences that flank the donor integration site. We show that testing a small number of sgRNAs, spanning multiple introns, is sufficient to identify a successful tagging site, and since these do not require a loci-specific donor, costs are minimal. While more information is needed to better understand how to integrate tags within proteins in the least disruptive way, this approach simplifies the generation of knock-in cell lines and makes scalable gene tagging highly accessible.

## Methods

### Cloning

The mNG2_11_ donor tag^4^ flanked by flexible 15 amino acid linkers was synthesized as two complementary oligos from IDT and annealed. This template was amplified by primers to add splice donor and acceptor sites, sgRNA target sequences external to the splice sites, and 25 nucleotide overhangs into the pMC.BESPX-MCS2 parental vector (System Biosciences). pMC.BESPX-MCS2 was digested with EcoRI and ApaI, and combined with the mNG2_11_ amplicon by Gibson assembly (NEB), generating the pMC-mNG2_11_ donor plasmid (Supplementary Data 1). The pMC-mClover3 donor plasmid was generated by replacing the mNG2_11_ sequence from the pMC-mNG2_11_ plasmid with the sequence of mClover3 (Addgene #74257) by Gibson assembly (Supplementary Data 1).

To generate HEK293 cells stably expressing mNG2_1-10_, mNG2_1-10_^4^ fused to the self-cleaving 2A peptide and tdTomato (Addgene #37347) was cloned into the lenti dCAS-VP64_Blast (Addgene #61425) backbone in place of dCas9-VP64 by 3-piece Gibson assembly.

sgRNA-expressing plasmids (Supplementary Table 1) were generated by digesting a lentiGuide-Puro plasmid (Addgene #52963) with Esp31 and ligating an annealed sgRNA oligo duplex as described previously^3^.

### Cell culture and transfection

Transfection experiments were carried out in HEK293 (ATCC CRL-1573) and HeLa cells (ATCC CCL-2). The HEK293 cells were generated to constitutively express mNG2_1-10_ and tdTomato from a stably integrated lentiviral cassette. Individual clones were sorted based on the tdTomato signal and a line with stable expression over time was selected for experiments.

Cells were cultured in DMEM (Thermo Fischer Scientific) + 10% fetal bovine serum (FBS; VWR) + antibiotic-antimycotic (Thermo Fisher Scientific). 4 × 10^6^ cells were plated across a 12-well plate the day prior to transfection. The donor plasmid was delivered at 5x the molar ratio of lentiCas9-Blast plasmid (Addgene #53962) and the two lentiGuide-Puro plasmids (Addgene #52963) encoding (1) the donor-cutting sgRNA and (2) the genomic locus-targeting sgRNA (Supplementary Table 1). In total, ∼1.4 µg of DNA were delivered to each well in 100 µL optimem (Thermo Fischer Scientific) with 4.3 µL of 1g/L PEI (Polysciences, cat. #24765). The following day, each well was expanded into a 10 cm dish. After six days, cells were harvested, analyzed by flow cytometry, and subjected to genomic DNA extraction.

### Flow cytometry and cell sorting

Cultured cells were trypsinized, resuspended in DMEM to ∼1 × 10^6^ cells/ml, and filtered through a cell strainer. Cellular fluorescence was measured on a BD FACSAria Fusion (BD Biosciences). Polyclonal fluorescent cell populations were acquired by isolating 1000 cells by sorting. Data were analyzed using Flowing Software 2 ver. 2.5.1 (http://flowingsoftware.btk.fi/index.php).

### Confocal microscopy and image processing

For imaging experiments cells were grown on coverslips and directly fixed in 4% formaldehyde (Electron Microscopy Sciences) in PBS (Thermo Fischer Scientific). Fixed cells were washed in PBS and coverslips were mounted on microscopy slides in Vectashield mounting medium (Vector Laboratories). Images were acquired on a Leica TCS SP8 confocal microscope. Z-stacks (0.6 μm slices) spanning the entire volume of the cells were recorded with oil-immersion 63× Plan-Apochromat lenses, 1.4 NA. Images were processed using Fiji^17^.

### Target analysis

Roughly 2 to 3 × 10^6^ cells were harvested for genomic DNA extraction in 100 μL of QuickExtract (Epicentre) according to manufacturer’s protocol. Amplification of edited genomic regions was performed with the EmeraldAmp MAX PCR Master Mix (Takara Bio USA). Primers were designed using the default parameters of primer3 (http://primer3.ut.ee/) to produce amplicons 250-300 nucleotides in length at the 5’ and 3’ junctions of each targeted site. Amplification reactions included a genomic primer upstream of the target integration site paired with a reverse primer hybridizing to the 3’ end of the tag, or a genomic primer downstream of the target integration site with a forward primer hybridizing to the 5’ end of the tag. The amplicons were imaged and extracted from a 2% agarose gel using the Monarch Gel Extraction kit (New England Biolabs), and analyzed by Sanger sequencing (GENEWIZ) using the tag-hybridizing primers from the amplification reaction.

